# p300 Is an Obligate Integrator of Combinatorial Transcription Factor Inputs

**DOI:** 10.1101/2023.05.18.541220

**Authors:** John J. Ferrie, Jonathan P. Karr, Thomas G.W. Graham, Gina M. Dailey, Gloria Zhang, Robert Tjian, Xavier Darzacq

## Abstract

Transcription coactivators are proteins or protein complexes that mediate transcription factor (TF) function. However, they lack DNA binding capacity, prompting the question of how they engage target loci. Three non-exclusive hypotheses have been posited: coactivators are recruited by complexing with TFs, by binding histones through epigenetic reader domains, or by partitioning into phase-separated compartments through their extensive intrinsically disordered regions (IDRs). Using p300 as a prototypical coactivator, we systematically mutated its annotated domains and show by single-molecule tracking in live cells that coactivator– chromatin binding depends entirely on combinatorial binding of multiple TF-interaction domains. Furthermore, we demonstrate that acetyltransferase activity negatively impacts p300–chromatin association and that the N-terminal TF-interaction domains regulate that activity. Single TF-interaction domains are insufficient for both chromatin binding and regulation of catalytic activity, implying a principle that could broadly inform eukaryotic gene regulation: a TF must act in coordination with other TFs to recruit coactivator activity.

## Introduction

Eukaryotic transcription factors (TFs) depend on coactivators to activate transcription. Coactivators have no intrinsic ability to bind specific DNA sequences, and their recruitment to specific target loci is instead proposed to rely on interactions with TF activation domains, modified histone tails, and other coactivators. Accordingly, most coactivators are multi-domain or multi-subunit complexes containing both enzymatic domains and diverse protein-protein interaction modules, including epigenetic reader domains (e.g., bromodomain, PHD domain), TF-interaction domains (e.g., KIX, PAS), and intrinsically disordered regions (IDRs).^4–6^ The relative importance of these various domains for target binding remains unclear. Moreover, it is puzzling how coactivators get distributed among target loci rapidly and reproducibly despite being stoichiometrically limiting relative to TFs and even to active cis-regulatory elements.^1–3^

To clarify these outstanding uncertainties in the field, we focused on p300—a central node in gene regulation that combines many domains in one polypeptide, making it more genetically tractable than multi-subunit coactivator complexes (e.g., Mediator). p300 is composed of three large regions: the N-terminal region (NTR), a well-structured enzymatic central region (Core), and the C-terminal region (CTR) (Figure 1A). In the Core are several annotated chromatin-interaction domains (ChIDs), while interspersed through the highly disordered NTR and CTR are various TF-interaction domains (TFIDs)—small helical and zinc finger domains that have been shown to engage in “fuzzy” binding with TF activation domains.^7^

**Figure 1.**
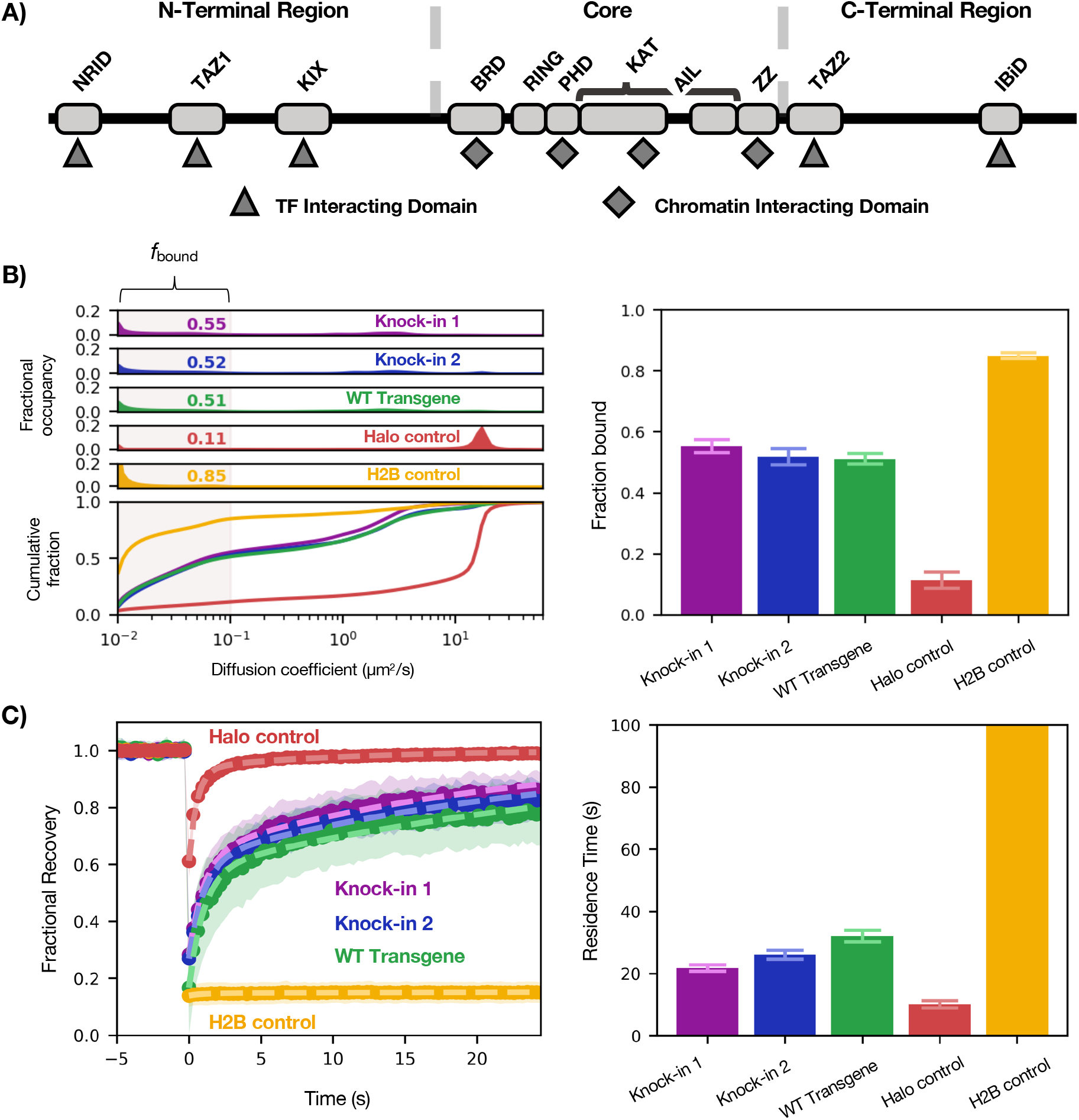
Benchmarking Halo-p300 diffusive behavior. (A) Schematic of domain organization of p300 showing the TFIDs in the NTR and CTR and ChIDs in the Core. The black line indicates IDRs. Relative lengths are not to scale. (B) Diffusive spectra (left)—probability density function (top) and cumulative distribution function (bottom)—with plot of *f_bound_* (right). Bars represent bootstrapping mean ± SD. (C) FRAP curves of the same (left) with residence times plotted for each construct (right). Bars represent best fit ± 95% CI.

We set out to address two questions: First, do ChIDs, TFIDs, or IDRs—or some combination thereof—mediate p300 chromatin engagement? Our hypothesis was that the TFIDs provide sequence specificity to direct p300 to target sites, while the ChIDs bind to histones to stabilize p300 on chromatin. By recognizing particular histone marks, the ChIDs could also contribute to directing p300 to target sites bearing active marks.^8^ Others have also argued that the IDRs of p300 cause it to partition into TF condensates.^9^ The second question we asked was whether there is interplay between p300 chromatin binding and catalytic activity. Because the p300 Core has both an acetyltransferase domain and a bromodomain—which is thought to bind acetylated lysines, especially on histones^10^—we considered that enzymatic activity might enhance chromatin binding and stabilize active p300 at its target sites.

## Results

To address these questions, we turned to high-speed (4 ms/frame) single-molecule tracking (SMT) combined with a Bayesian analysis method that infers underlying diffusive states from a population of observed trajectories.^11^ The output of this analysis is a “diffusion spectrum,” or probability density for every value of diffusion coefficient (Figure 1B). The fractional likelihood of a molecule moving with a diffusion coefficient of <0.1 μm^2^s^-1^ we call the bound fraction (f_bound_) because it represents the proportion of molecules diffusing at a rate indistinguishable from that of chromatin motion (see Halo-H2B in Figure 1B). By measuring their effects on f_bound_, we are thus able to assess the impact of various p300 mutations and perturbations on chromatin interactions in the context of the live cell.^11, 12^ Although the rest of the diffusion spectrum faster than 0.1 μm^2^s^-1^ holds other valuable information, such as how many distinct diffusive species (number of peaks) exist for a given tagged protein, we have mainly focused on changes to f_bound_ in this report.

We first benchmarked SMT behavior of the gene products of two HaloTag knock-ins at the endogenous EP300 locus against stably integrated Halo-H2B and Halo-NLS transgenes in U2OS cells (Figure 1B-C). As expected, Halo-H2B was predominantly bound (f_bound_ = 0.85 ± 0.01) while Halo-NLS was predominantly fast-diffusing. (We note that Halo-NLS has an f_bound_ of 0.11 ± 0.03, which we consider the baseline of the assay.) In this cell line, p300 had an f_bound_ of approximately 0.54, which was reproducible between two different clonal knock-in lines (f_bound_ = 0.55 ± 0.02, 0.52 ± 0.03). We also used FRAP to measure the residence time of each p300 construct on chromatin and found that in contrast to H2B, which has a residence time far beyond the timescale of this experiment, p300–chromatin binding events persist for approximately 26 seconds on average—on par with or slightly longer than residence times typical of TFs.^13^

Having established a characteristic profile for WT endogenous p300 diffusion, we built a transgene system to facilitate mutation of p300. In order to avoid complexities arising from interactions with endogenous p300, we generated a clonal knock-out cell line expressing no detectable p300 (Fig. S1A) into which various p300 transgenes were introduced by random integration and antibiotic selection. The full-length p300 transgene product behaved similarly to the tagged endogenous protein (f_bound_ = 0.51 ± 0.02—Figure 1B-C), validating this assay system. The stable transgene was considerably less expressed than the endogenous protein, but we verified that in our system f_bound_ is not sensitive to concentration of protein (Fig. S2).

### p300 Core is neither necessary nor sufficient for chromatin binding

To address which domains mediate p300 chromatin engagement, we first assessed which of its three regions (NTR, Core, CTR) is sufficient for binding (Figure 2B). Strikingly, the Core had essentially no ability to bind on its own (f_bound_ = 0.09 ± 0.01). The NTR exhibited a modest capacity to bind (f_bound_ = 0.22 ± 0.02), while CTR was sufficient to reach full-length p300 levels of binding (f_bound_ = 0.50 ± 0.02). Both NTR and Core constructs had significantly reduced residence times (t = 16, 14 s) compared to CTR and p300 (t = 26, 26 s) (Fig. S3).

**Figure 2.**
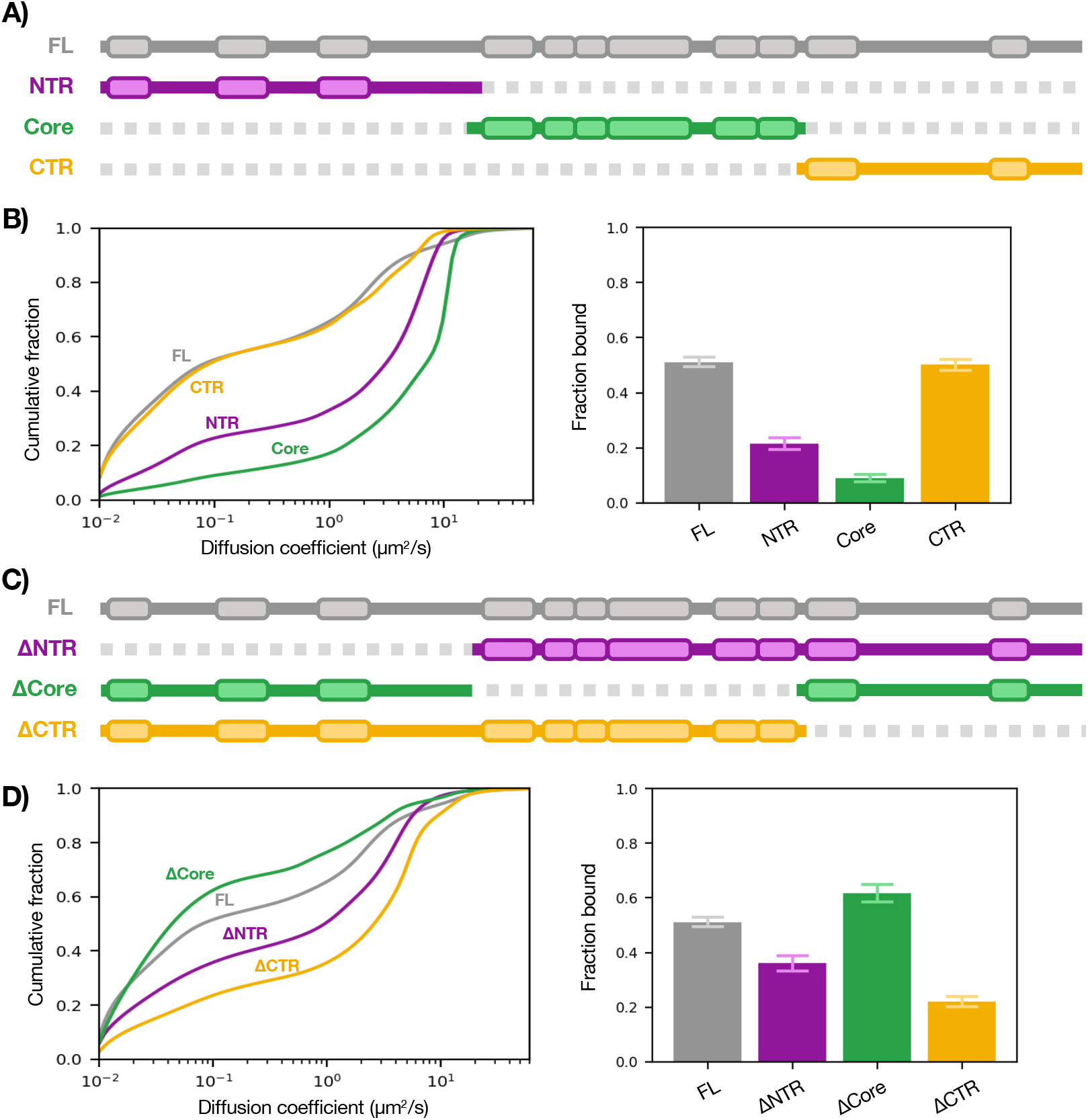
Sufficiency and necessity of p300 regions for chromatin binding. (A) Schematic of p300 regions. (B) SMT of isolated regions of p300 (left) with summary bar plot of *f_bound_* (right). (C) Schematic of p300 truncations. (D) SMT of truncations of p300 (left) with summary bar plot of *f_bound_* (right). Bars represent bootstrapping mean ± SD. See also Fig. S3.

We then asked which of the domains is necessary for binding by tracking truncations of p300: NTR-Core (ΔCTR), Core-CTR (ΔNTR), and NTR-CTR (ΔCore) (Figure 2D). Remarkably, ΔCore had a somewhat enhanced f_bound_ of 0.62 ± 0.03—a surprising result we return to later. ΔNTR and ΔCTR showed significantly decreased ability to bind chromatin (f_bound_ = 0.36 ± 0.03, 0.22 ± 0.02) compared to WT p300 (f_bound_ = 0.51 ± 0.02). Together, these results suggest that the ChID-containing Core is dispensable for p300–chromatin binding in vivo while the NTR and CTR are necessary and sufficient (albeit to different extents). To confirm this, we performed SMT on a series of Core mutants which had affected p300 binding and activity in vitro (including a bromodomain mutant) and saw no substantial changes in our in vivo assay (Fig. S4).

### p300 TFIDs are necessary for chromatin binding

The finding that p300’s chromatin-binding capacity lies outside its Core domain does not necessarily mean that the TFIDs are what mediate binding—it could be the IDRs, which compose the vast majority of the NTR and CTR. (Whereas each TFID is 50-80 aa, there is approximately 1,400 aa of IDR.) We therefore measured the sufficiency of the TFIDs for chromatin binding by expressing them as HaloTag fusions and observed that each TFID has only a modest capacity to bind when acting alone (Figure 3B). Next, we tested the necessity of these domains by deleting each of the TFIDs as well as all the TFIDs in the otherwise full-length protein and performed SMT (Figure 3D). Although each TFID deletion only partially impaired chromatin binding, when all TFIDs were deleted (ΔALL) there was a drastic reduction in f_bound_ to just above the baseline (0.14 ± 0.02). Hence, it appears that the combined action of multiple TFIDs is required to bring p300 to chromatin. The finding that deleting all TFIDs essentially incapacitated chromatin binding also suggests that the IDRs are not sufficient for p300– chromatin association. Additionally, the lack of binding by the ΔALL construct is another indication that the Core is not sufficient for p300-chromatin binding.

**Figure 3.**
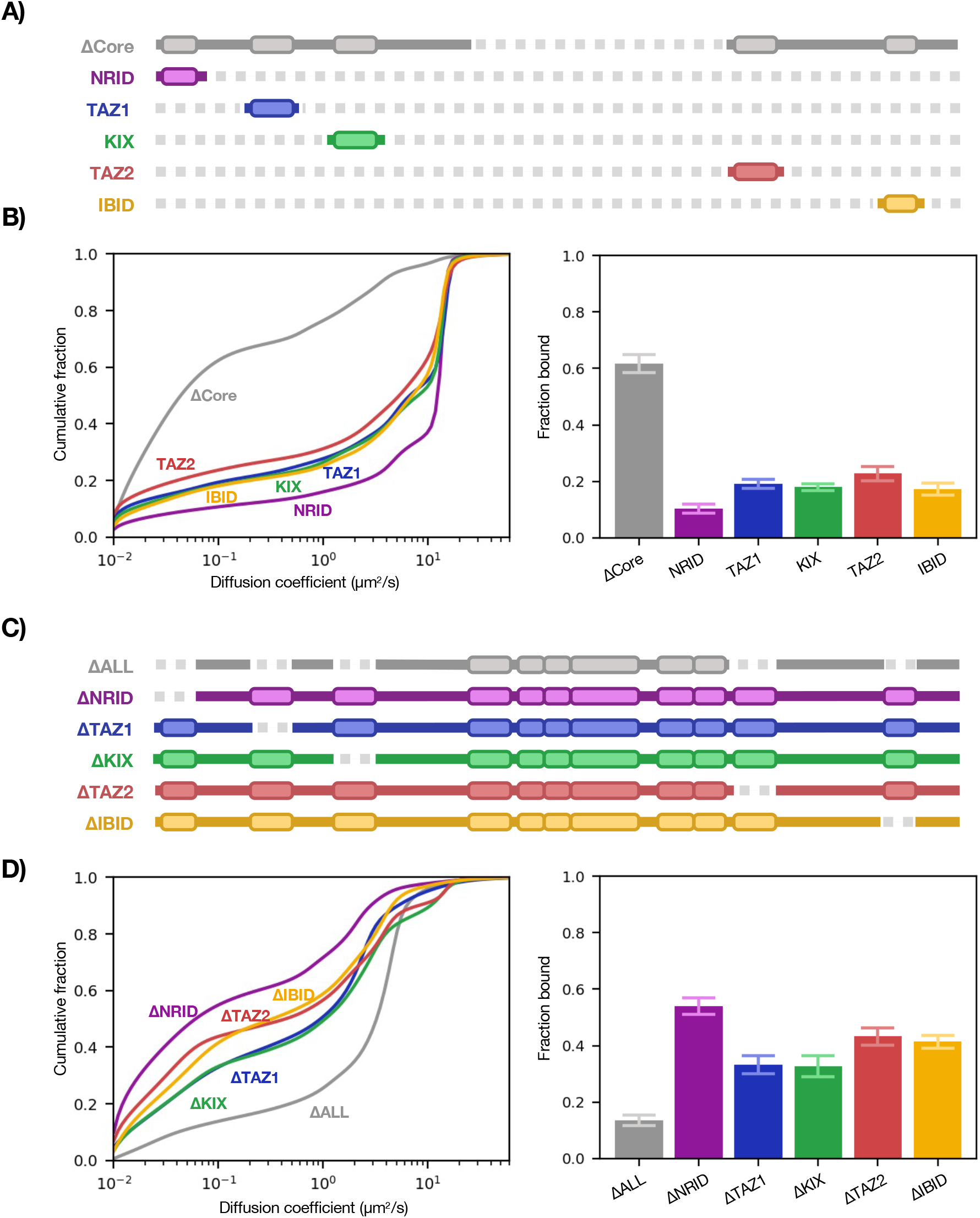
Sufficiency and necessity of p300 TFIDs for chromatin binding. (A) Schematic of p300 TF interaction domains (core region in grey). (B) SMT of TFIDs (left) with summary bar plot of *f_bound_* (right). (C) Schematic of p300 TFID deletions. (D) SMT of TFID deletions of p300 (left) with summary bar plot of *f_bound_* (right). Bars represent bootstrapping mean ± SD.

### Acetyltransferase activity opposes p300–chromatin binding

Although it seems that the Core does not contribute appreciably to p300–chromatin binding, it is intriguing that its deletion increased p300 f_bound_ from 0.51 ± 0.02 to 0.62 ± 0.03. We wondered whether this was the consequence of the loss of its acetyltransferase activity and tested this in two ways: by imaging a catalytically dead mutant (Y1467F)^14^ and tracking WT p300 after addition of the potent and specific catalytic inhibitor A485.^15^ Both perturbations increased p300 f_bound_ similar to complete loss of the Core domain (Figure 4A), which tallies with previous ChIP-seq data showing an increase of p300 peak heights upon A485 treatment.^16^ To demonstrate that the response to A485 was a direct effect of p300 activity and not an indirect consequence of some off-target cellular response, we designed a p300 point mutant that remains catalytically active in the presence of A485 (H1451K) (manuscript in preparation) and as expected, saw no change by FRAP (Figure 4B) or SMT (Fig. S5) upon A485 addition.

**Figure 4.**
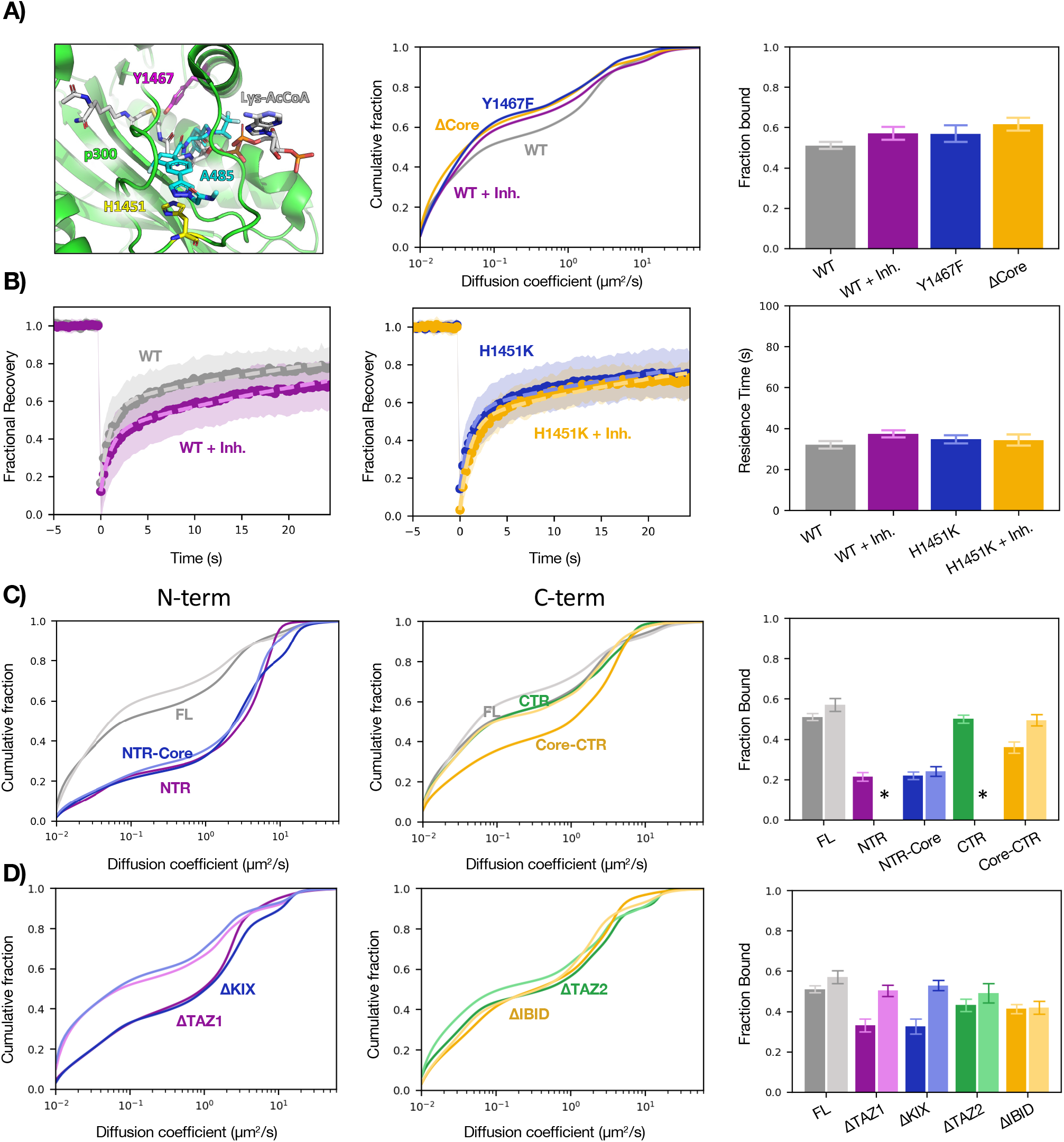
Effects of the catalytic core on p300–chromatin interactions. (A) Cartoon representation (left) of p300 active site showing residues H1451 (pink) and Y1467 (yellow), substrate mimetic (grey), and A485 inhibitor (teal). Plots comparing diffusion spectra (center) of WT p300 and three perturbations of p300 catalytic activity: addition of inhibitor A485, active site mutation Y1467F, and deletion of the catalytic core, with bar plot of *f_bound_* (right). (B) FRAP plots of WT (left) and mutant (center) in response to A485 with residence times plotted for each construct (right). (C) SMT of NTR-containing (left) and CTR-containing (center) constructs in the presence (lighter hue) or absence (darker hue) of A485, with bar plot of *f_bound_* (right). *Data not acquired. (D) SMT of N-terminal ΔTFID (left) and C-terminal ΔTFID (center) constructs in the presence (lighter hue) or absence (darker hue) of A485, with bar plot of *f_bound_* (right). SMT: bars represent bootstrapping mean ± SD. FRAP: Bars represent best fit ± 95% CI. See also Fig. S5.

The finding that catalytic activity opposes chromatin binding provided a possible explanation for two curious observations: ΔNTR (Core-CTR) had reduced binding compared to CTR and deletion of the two N-terminal TFIDs had a greater impact on f_bound_ than deletion of the C-terminal TFIDs—both of which are strange because the CTR is both necessary and sufficient for full chromatin association. We therefore reasoned that the N-terminus may function (directly or indirectly) to inhibit Core catalytic activity and thereby counteract the Core’s effect to reduce CTR binding. Indeed, the reduction in f_bound_ from adding the Core to the CTR could be rescued by treatment with A485 (Fig. 4C). Furthermore, ΔTAZ1 and ΔKIX were both hyper-sensitive to the drug compared to the FL with respect to f_bound_ (Fig. 4D), indicating that the N-terminal TFIDs participate in negative regulation of core catalytic activity.

## Discussion

Combining high-speed SMT with the domain-mapping strategy classically used in in vitro biochemistry allowed us to determine which domains of a modular coactivator determine its chromatin binding in living cells. The results indicate that p300 binds chromatin primarily through multivalent TF-TFID interactions, not by its ChIDs or IDRs (Figure 5A).

**Figure 5.**
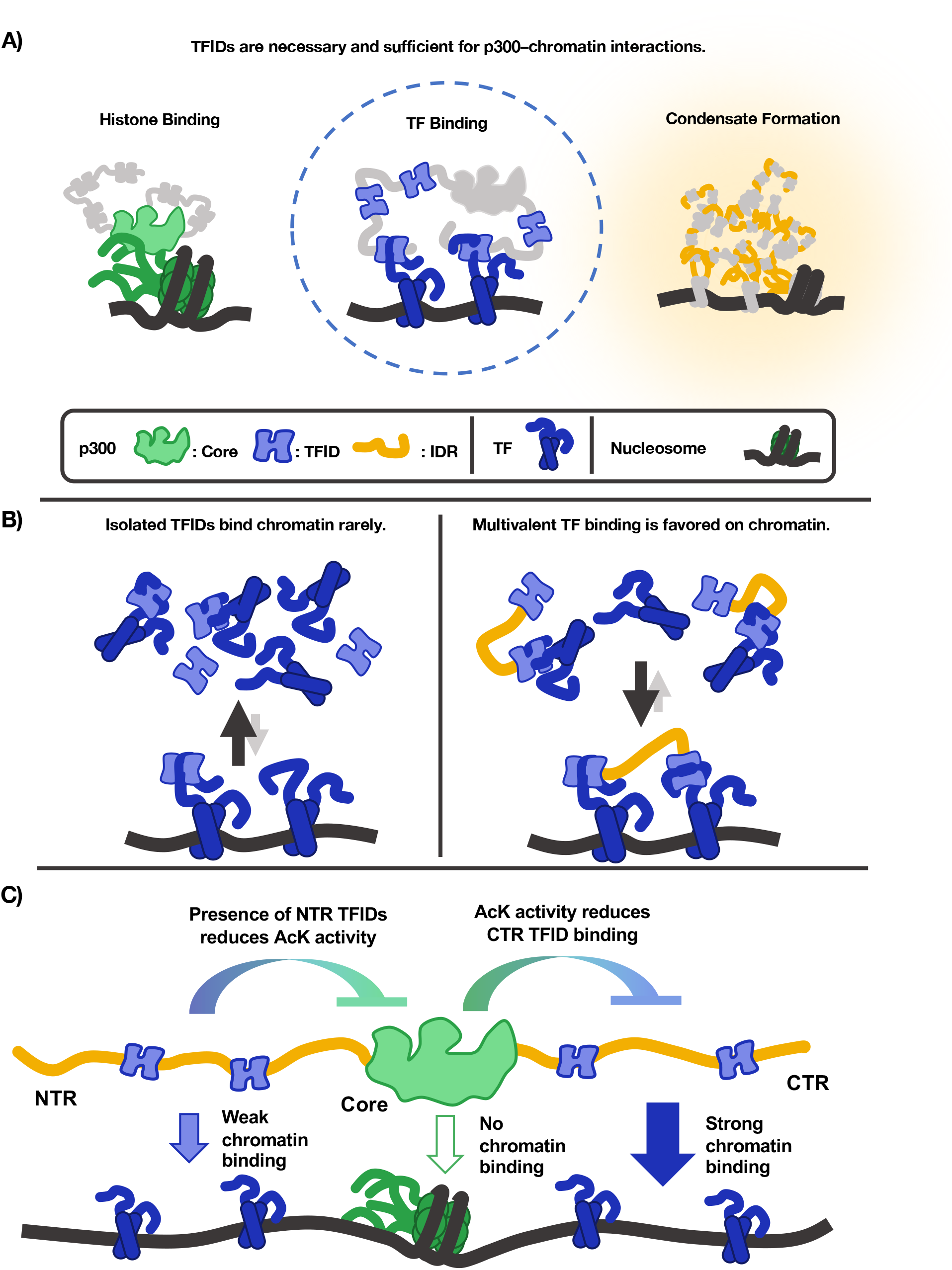
Graphical model of p300–chromatin interactions. (A) Three existing models of how p300 engages with chromatin, the second of which is strongly supported by in vivo SMT data. (B) Multivalent TFID-TF interactions favor the chromatin-bound state. (C) The contributions of each p300 domain to p300–chromatin interactions.

### SMT discriminates between models of p300–chromatin engagement

Two measurements we made are particularly clarifying: Deleting the Core increased f_bound_ to 0.62, while removal of the five small TFIDs essentially eliminated p300 chromatin binding, bringing f_bound_ down to 0.14 (Figure 3). Thus, the collective action of the small helical peptides we call TFIDs is both necessary and sufficient for p300–chromatin interactions. While the Core and the IDRs undoubtedly contribute to other functions, our data consistently show that the way p300 associates with chromatin is not through the Core domains binding histones or IDR-mediated interactions but through TFID-mediated interactions with sequence-specific TFs. (Though it is possible, of course, that factors other than sequence-specific TFs also engage p300 TFIDs.) These findings comport with in vitro binding assays showing that bromodomains have micromolar affinity for acetylated histone peptides^10^ while TFIDs have nanomolar affinity for TF peptides.^17, 18^ Furthermore, the increased f of ΔCore relative to WT suggests that catalytic activity does not cause p300–chromatin association through its bromodomain, which is confirmed by the finding that chromatin binding ability was not impaired by the acetyl-transferase inhibitor A485, a catalytically dead mutant, or the p300 bromodomain mutant (Fig. 4B, Fig. S4).

### Multiple TFID interactions enable avid-like binding of p300 to chromatin

Beyond addressing our initial questions, our investigation yielded insights into the mechanism of p300 recruitment. The finding that each TFID on its own largely occupies the unbound state suggests that the majority of each individual TFID’s binding partners are unbound, and that a single TFID has little or no preference for chromatin-bound TFs. (We consider it unlikely that the TFID is merely unbound by a TF, given the multiplicity of TF binding partners and the aforementioned nanomolar affinities measured between TFs and p300 TFIDs.) Multimerization of TFIDs can strongly favor interactions with chromatin, as demonstrated by the fact that a two-TFID fragment (CTR) has a dramatically greater chromatin-bound fraction than either TFID alone. A simple explanation for this is an avidity effect of engaging multiple TFs bound to adjacent DNA sites. Binding multiple TFs on chromatin is energetically more favorable than binding them in solution, as the entropic cost of bringing the TFs together is already paid. In principle, a coactivator with multiple TFIDs could even have a higher chromatin-bound fraction than any of the TFs it binds. (For why we disfavor the hypothesis that additional TFIDs increase p300 f_bound_ by increasing the number of possible binding sites on chromatin, see Supplemental Note.)

### p300 is an obligate integrator of combinatorial TF inputs

The requisite for multiple TFIDs in p300 recruitment suggests that p300 functions as an obligate integrator—that is, it not only can but must bind multiple TFs to associate with chromatin. Such integration has profound implications for p300 recruitment at two levels—in trans and in cis. In trans, any gene-regulatory process that requires the action of p300 must deploy multiple TFs that can engage multiple TFIDs. In cis, any regulatory element that requires p300 activity must be able to bind more than one of its TF partners, rendering isolated TF binding sequences insufficient for recruiting p300 in a competitive environment. If this is a general rule for coactivator recruitment, the high degeneracy of eukaryotic TF recognition motifs^19^ might be straightforwardly rationalized: Because a TF must act through recruiting coactivators in coordination with another TF, their modest individual specificities are multiplied to recruit coactivators to cis regulatory elements that are capable of binding both TFs simultaneously—thus solving the problem we posed of how a limiting pool of coactivators is distributed among target loci. Those factors that can engage more TFIDs on their own, such as p53,^20^ may be the more context-independent activators, while those that can only engage one of the TFIDs are likely more dependent on other proteins (i.e., additional TFs) to achieve activation of target genes.

Last, because p300 binding is sensitive to catalytic activity, our assay was able to uncover a new mode of p300 catalytic regulation by the N-terminal TFIDs (Fig. 4C-D). The Core is most active in the absence of TAZ1 and KIX, which suggests that these domains or their binding partners somehow modulate acetyltransferase activity, which activity inhibits the chromatin binding of the C-terminal TFIDs (Fig. 5C). While these data do not shed light on the mechanism of such regulation, there is an important biological consequence of the observation: That p300 catalytic activity is regulable through two of its TFIDs shows another way in which it is an obligate integrator of TF inputs.

## Acknowledgments

We would like to thank all the members of the Tjian-Darzacq lab for many rich discussions and helpful suggestions over the years, and Vinson Fan in particular for his advice on data analysis. We also thank Luke Lavis for providing fluorescent HaloTag ligands and the CRL Flow Cytometry Facility for use of their instruments. This work was supported by National Institutes of Health grants U54-CA231641-01659 (to XD), 5T32GM007232-42 (supporting JPK), the Silicon Valley Community Foundation/CZI, and the Howard Hughes Medical Institute. JJF is a Howard Hughes Medical Institute Awardee of the Life Sciences Research Foundation.

## Author Contributions

JJF and JPK devised and executed experiments with support from GZ and GD under the guidance of RT and XD. JJF and JPK analyzed the data with help from TWG. JPK drafted the main text of the original manuscript. XD and RT supervised the project.

## METHODS

### RESOURCE AVAILABILITY

#### EXPERIMENTAL MODEL

Human U-2 osteosarcoma cells (U2OS) (RRID: CVCL_0042) were a gift from David Spector’s lab, Cold Spring Harbor Laboratory. Cells were cultured at 37°C with 5% CO_2_ in homemade phenol red containing DMEM (Thermo Fisher #12800082) with 4.5 g/L glucose supplemented with 10% fetal bovine serum (HyClone, Logan, UT, Cat. #SH30396.03, lot #AE28209315), 1 mM sodium pyruvate (Thermo Fisher 11360070), L-glutamine (Sigma # G3126-100G), GlutaMax (Thermo Fisher #35050061), and 100 U/mL penicillin-streptomycin (Thermo Fisher #15140122). Cells were subcultured at a ratio of 1:4-1:10 every 2-4 days. Regular mycoplasma testing was performed by PCR. Phenol red-free DMEM (Thermo Fisher #21063029) supplemented with 10% fetal bovine serum, 1 mM sodium pyruvate (Lonza 13-115E), and 100 U/mL penicillin-streptomycin was used for imaging.

### METHOD DETAILS

#### Cell culture and stable cell line construction

Stable cells lines expressing exogenous gene products (Table 1) were generated by PiggyBac transposition and antibiotic selection. Gibson Assembly was used to clone full-length p300 and p300 fragments/mutants into a PiggyBac vector containing a puromycin resistant gene and 3X-FLAG-Halo upstream of a multiple cloning site. All cloning was confirmed by Sanger sequencing. Cells were lipofected using Lipofectamine 3000 (Thermo Fisher #L3000015). Following manufacturer instructions, 125 uL of Opti-MEM medium (Thermo Fisher #31985062) was combined with 5 uL of P3000 Reagent, 3-5 ug of gene containing PiggyBac vector, and 1.5 ug of SuperPiggyBac transposase vector which was subsequently mixed with 125 uL of Opti-MEM Medium following addition of 5 uL of Lipofectamine 3000 Reagent. After brief vortexing the combined media was added directly to a 6-well containing 70-90% confluent cells. Transfected cells were cultured for 1-2 days before returning the cells to normal DMEM growth media. Once confluent, cells were passaged 1:8 and placed in selection media—DMEM growth media containing 1 ug/mL puromycin (Thermo Fisher #A11138-03)—and selected for 7 days. Polyclonal stable cell lines were maintained in selection media following selection.

#### CRISPR/Cas9-mediated genome editing

Knock-in and knock-out cell lines were created as published previously.^21^ We transfected U2OS cells using Lipofectamine 3000 (ThermoFisher L3000015) according to the manufacturer’s protocol. For knock-out generation, 1 μg of each Cas9 vector was transfected per well in a 6-well plate; for knock-in generation, 2 μg repair vector and 1 μg Cas9 vector per well in a 6-well plate.

For Halo-p300 knock-in, sgRNA sequences were designed with the help of the CRISPOR web tool to find and evaluate guides in 200bp window centered on insertion site. Guides with high specificity close to site of insertion were chosen. Repair vectors were cloned into basic backbone (pENTR) with tag with 500bp left and right homology arms. PAMs for the guides were mutated in pHDR by site-directed mutagenesis. sgRNAs were cloned into the Cas9 plasmid (a gift from Frank Xie) under the U6 promoter (Zhang Lab) with an mVenus reporter gene under the PGK promoter.^22^ Two guide/repair-vector pairs were attempted at each terminus; only N-terminal clones were ultimately recovered. 24-48 hours after transfection, a pool of mVenus positive cells were FACS-sorted and expanded for approximately a week, after which single Halo-positive cells (stained with JF549) were sorted into 96-well plates. Clones were expanded for ∼3 weeks and genotyped with immunostaining to confirm.

For p300 knock-out, sgRNA sequences from knock-ins were used in combination—one from either terminus of EP300. 24-48 hours after transfection, single mVenus positive cells were FACS-sorted into 96-well plates. Clones were expanded for ∼3 weeks and then genotyped with immunostaining to confirm.

#### Cell preparation and dye labeling for imaging

For both SMT and FRAP, ∼200,000 cells were plated in a 35 mm dish containing either a 20 mm diameter or 14 mm wide uncoated coverslip (MatTek Corporation #P35G-1.5-20-C or #P35G-1.5-14-C) one day prior to imaging. Before imaging cells were labeled with either HaloTag-JF549 or JFX549 dye at 5 nM, for SMT, or 50 nM, for FRAP. After 20 min of incubation, free dye was removed by replacing media with 2 mL of phenol-red free media and incubating for 15 minutes. Following a second media replacement and incubation, the media was replaced a final time and cells were imaged.

#### Flow cytometry

Cells were grown in 12-well dishes, labeled with ∼50 nM JF549 for 45 minutes, and washed three times in dye-free DMEM fifteen minutes each before trypsinization, filtration, and flow. Used PE filter on a BD Bioscience LSR Fortessa.

#### Antibodies

Endogenous p300 was detected using Abcam 48343, while transgenes were detected using Millipore Sigma F3165 (anti-FLAG)—both of which were followed by Invitrogen 31430 (anti-mouse-HRP).

#### Western blotting

Samples for Western blots were prepared by either nuclear extraction or direct lysis. Nuclear extraction was performed by resuspending ∼8 million cells in 100 uL hypotonic buffer (100 mM NaCl, 25 mM HEPES, 1 mM MgCl_2_, 0.2 mM EDTA, 0.5% NP-40) containing protease inhibitors (Roche) on ice with gentle pipetting. Following resuspension, nuclei were isolated by centrifugation (5 min, 4000 g). Subsequently, nuclei were resuspended in 100 uL hypotonic buffer containing protease inhibitors and benzonase (THERMO, 0.2 uL per 100 uL) to digest DNA and incubated at 25 C with gentle agitation for 20 mins. Following incubation 4X LDS (16% BME, 200 mM Tris-HCl, 8% SDS, 40% glycerol, 0.4% bromophenol blue) was added to each sample and samples were boiled for >5 mins. Direct lysis was performed by pelleting ∼1E6 cells, resuspending cell pellets in 100 uL of LDS, and boiling for > 5 mins. Constructs which showed a high fraction bound (WT, NTR, CTR, ΔCore, ΔNRID, ΔTAZ1, ΔKIX, ΔTAZ2, ΔIBID, 1132, 1451, 1467, AIL, 1645) were prepared as nuclear extracts as they could not be visualized using direct lysis. Wet transfer was performed overnight at constant current (90 mA) in transfer buffer (15 mM Tris-HCl, 20 mM glycine, 20% methanol, 0.0375% SDS). Following Ponceau staining, membrane was blocked with 10% non-fat milk in TBS for 1 hour at room temperature with agitation. After removal of blocking solution, 5% non-fat milk in TBS containing primary antibody (∼20 ug) was added and incubated at room temperature with agitation for at least 1 hour. Membrane was briefly washed 3 times with TBS and 5% non-fat milk in TBS containing HRP-conjugated secondary antibody (1:5000) was added to the membrane and incubated at room temperature for 1 hour with agitation. Finally, secondary antibody containing solution was removed, and membrane was washed 3 time with TBST with short (∼2 min) incubations with agitation. Western blots were visualized with SuperSignal ^TM^ West Femto (Thermo Scientific #34094).

#### Single-molecule tracking

All SMT experiments were carried out on a custom-built microscope as previously described.^21^ In brief, a Nikon TI microscope is equipped with a 100X/NA 1.49 oil-immersion TIRF objective, a motorized mirror, a perfect Focus system, an EM-CCD camera, and an incubation chamber maintained with humidified atmosphere with 5% CO_2_ at 37 °C. All microscope camera and hardware components were controlled through the NIS-Elements software (Nikon).

Dye-labeled cells were excited with 561 nm laser at 1100 mW (Genesis Coherent, Santa Clara, CA) and imaged through an Semrock 593/40 nm bandpass filter. All imaging was performed using highly inclined laminated optical sheet (HiLo) illumination (Tokunaga 2008). Low laser power (2-5 %) was used to locate and focus cell nuclei, and an ROI (region of interest) of 100X100 pixels was positioned such that as much of a single nucleus as possible was captured. Subsequently samples were pre-bleached at high laser power (100%) until a limited number (1-4) single particles were visible per frame. Movies were taken with 1 ms pulses of full power of 561 nm illumination with a camera exposure of 4 ms per frame for 5,000-10,000 frames. At least 10 (but often no more than 25) movies were collected for each sample as one tehcnical replicate on a given day. Two technical replicates on two separate days were collected to produce the reported results. As observed by others^23^ the variance within collected data is largely due to cell-to-cell variance, with day-to-day variance being relatively minimal.

#### SMT data processing

Raw SMT movies were processed using the open-source software package quot (https://github.com/alecheckert/quot)^11^ to extract single molecule trajectories. The quot package was run on each SMT movie collected using the following settings: [filter] start = 0; method = ‘identity’; chunk_size = 100; [detect] method = ‘llr’; k = 1.2; w = 18; t = 18; [localize] method = ‘ls_int_gaussian’; method = ‘ls_int_gaussian’; window_size = 9; sigma = 1.2; ridge = 0.0001; max_iter = 10; damp = 0.3; camera_gain = 109.0; camera_bg = 470.0; [track] method = ‘conservative’; frame_interval = 0.00448; search_radius = 1; max_blinks = 0; min_I0 = 0; scale = 7.0.

SMT trajectories were analyzed using the open-source software package SASPT (https://github.com/alecheckert/saspt).^11^ The first 200 frames of each movie were removed due to high localization density. To confirm that all movies were sufficiently sparse to avoid misconnections, all movies maintained a maximum number of localizations per frame of six (though most frames had 1 or less). The StateArray method was used with the following settings: likelihood_type = ‘rbme’; pixel_size_um = 0.16; frame_interval = 0.00448; focal_depth = 0.7; start_frame = 200; sample_size = 1000000; splitsize = 10.

#### Fluorescence Recovery After Photobleaching

FRAP experiments were conducted on a Zeiss LSM 900. Dye-labeled cells were excited with 561 nm laser with a 150 μM pinhole. Laser power (0.7 – 7.5%), detector gain (700 – 900 V), detector offset (-60 – -13), and digital gain (1.0 – 1.5%) was adjusted between samples and cells, due to high variations in expression level between p300 constructs. For each cell, the ROI was adjusted to contain the whole nucleus and the scan speed was set to the maximum allowable for the frame size. Movies were collected using the max scan speed for the given ROI with bidirectional scanning, an 8-bit depth, and 300 msec delay between frames for 100 frames resulting in a 30-second-long movie. Bleaching was performed on frame 21 of the movie, allowing the first 20 frames to be used to normalize the fractional recovery. At least 20 (but often no more than 35) movies were collected for each sample as one technical replicate on a given day. Two technical replicates on two separate days were collected to produce the final results.

### QUANTIFICATION AND STATISTICAL ANALYSIS

#### Statistical analysis of SMT

To derive a measure of error, we performed bootstrapping analysis on all SMT datasets. For each dataset, a random sample of size n, where n is the total number of cells in the dataset, was taken 100 times. The mean and standard deviation of SASPT done on the 100 samples are reported.

#### FRAP data processing

Cells whose fractional recovery after a single frame was 95% due to ineffective bleaching or whose recovery could not be delineated from noise were discarded from further analysis. Average fractional recovery curves and fits to both average and single cell curves were performed within the same MATLAB script. Fractional recovery curves were computed and fit for each cell and sample through the following procedure:

1. **Nuclear masking.** A 5-pixel width Gaussian filter is applied to smooth the image. Nuclear masking is performed by thresholding (70% of the maximum intensity).
2. **Masking the Bleach Spot**. The bleach spot mask is generated from the location and size information imported from the metadata within the image. Bleaching of H2B was used as a control to determine the actual bleach spot size relative to the size specified during data acquisition.
3. **Segmented Intensity Calculation**. The sum intensity was computed for each frame within the nucleus outside of the bleach spot (I_nucleus_) and within the bleach spot (I_bleach_). Additionally, the mean intensity outside of the nucleus (I_outside_) was computed for each frame.
4. **Background Subtraction**. The background for each frame was computed by multiplying the mean pixel intensity outside of the nucleus (I_outside_) by the number of pixels inside the nucleus and inside the bleach spot to obtain the background intensity inside the nucleus and inside the bleach spot. The resultant per frame background intensities were subtracted from the respective sum intensity values.
5. **Fractional Recovery**. The fractional recovery is computed as:

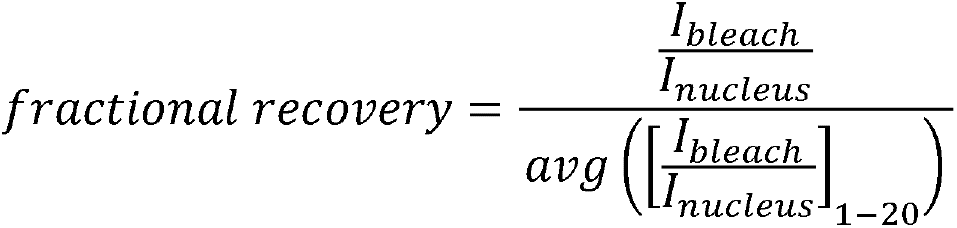 where the intensity ratio per frame is normalized by the average intensity ratio over the first 20 frames.
6. **Manual Curation.** Sum intensity projections of the first 20 frames of each acquired movie were inspected along with the intensity difference between the frames before and after bleaching that had been gaussian filtered in Fourier space. Cells that either appeared dead or showed other issues of cell health (nuclear blebbing, numerous large puncta, etc.) in sum intensity projections, who’s nuclear mask did not correctly contain nuclear intensity, or that had a significant portion of intensity differences outside of the bleach region were removed from analysis.
7. **Average Recovery Curves for Samples.** Due to the high degree of variability in signal intensity from cell to cell, which required optimization of the imaging conditions for each cell, the average recovery curve was computed as a weighted average of the single cell recovery curves based on their signal to noise. Curves were weighted by the inverse of the variance of the intensity of the first 20 frames of the movie, resulting in higher contributions to the average recovery from cells with higher signal to noise.
8. **Fitting Fractional Recovery Curves**. We assessed the impact of having a different bleach spot radius on the recovery profile of WT p300, prior to selecting a model for fitting the FRAP data. By comparing the FRAP curves using 2 and 3 uM bleach spot radii, we observed (Fig SX) that the recovery profile was not dependent on the bleach spot size indicating that we are not observing diffusion and are in a binding reaction dominant regime.^24^ Therefore, fractional recovery profiles were fit two two-exponential model:

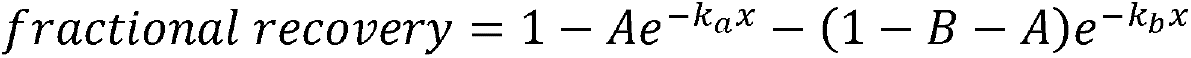 where A is the fraction of molecules that recover quickly, B is bleach depth (and therefore 1-B-A is the fraction of molecules that recover slowly), k_a_ is the recovery rate of molecules that recover quickly, and k_b_ is the recovery rate of molecules that recovery slowly. For all fits, the bleach depth (B) was fixed to the fractional recovery of the first frame following bleaching. All samples were initially fit allowing A, k_a_, and k_b_ to vary. Single exponential fits were attempted but were sufficiently poor to warrant addition of a second exponential recovery term. All fitting was performed using a weighted fitting algorithm where the weight of any given datapoint was equal to the absolute difference in intensity between a given frame and the subsequent frame. This biased the fitting towards the data rich, initial recovery portion of the fractional recovery curves. Subsequently, to further limiting overfitting, fitting for all p300 samples was constrained by fixing the fast recovery rate (k_a_) at 0.95 s^-1^ which was close the average observed value and was optimized to produce high quality fits of both relatively fast and slow recovering constructs (Table S3). Fit quality was nearly identical allowing only A and k_a_ to vary.

