## Supplemental information for "p300 Is an Obligate Integrator of Combinatorial Transcription Factor Inputs"

### SUPPLEMENTAL FIGURES

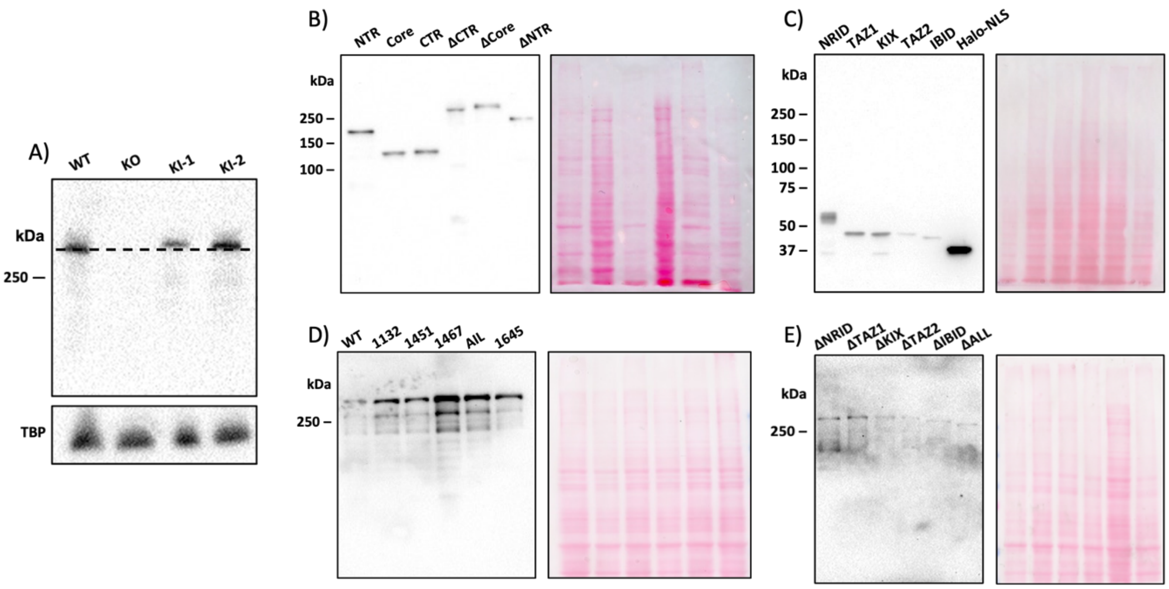


**Figure S1—Western blots of p300 variants.**
(A) Anti-p300 blot of unedited U2OS cells, the *EP300* knockout line, and the two Halo-p300 knock-in lines, with loading control beneath. Dashed line is to show shift in molecular weight from addition of HaloTag. (B-E) Anti-FLAG blots of all transgene constructs in the paper with Ponceau stains to the right as loading controls. Constructs were loaded to achieve comparable intensities—for relative expression levels, see Fig. S2.

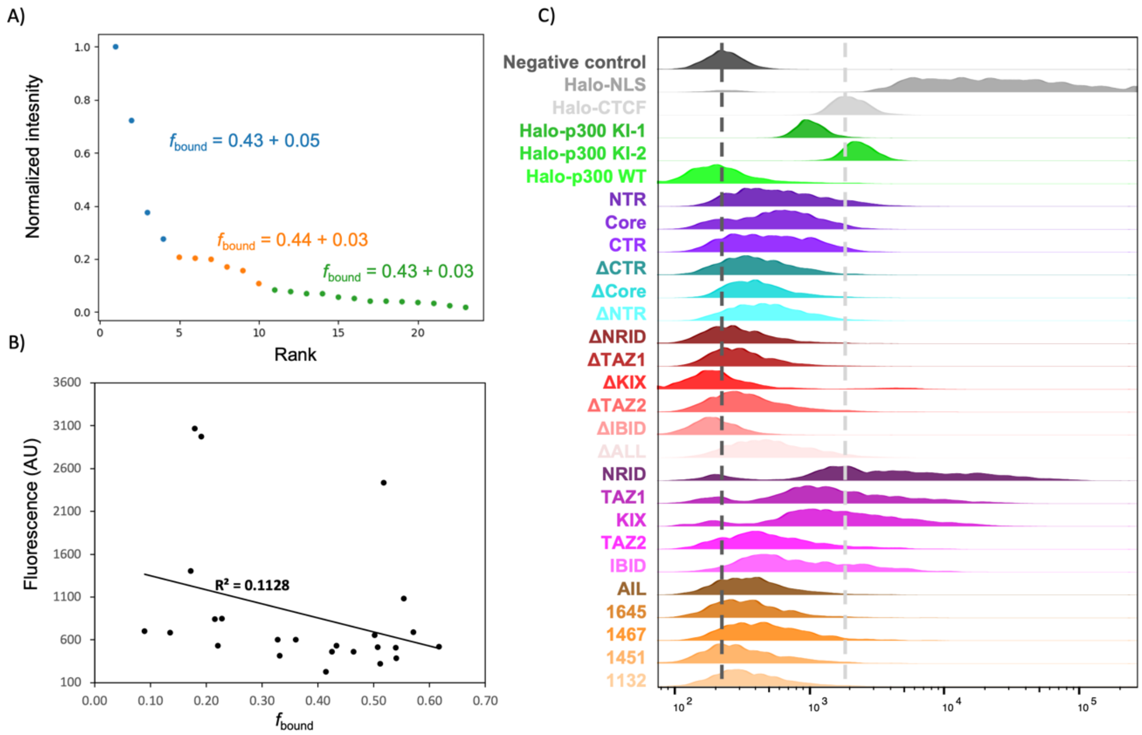


**Figure S2—Fraction bound is unaffected by p300 expression level.**

(A) Plot of cellular intensities from a transient transfection of WT p300 into *EP300* knockout cells with SMT results of three intensity bins overlaid. (B) All p300 mutants in the paper were measured by flow cytometry (C) and their mean cellular fluorescence plotted against SMT-derived $f_{\text{bound}}$.

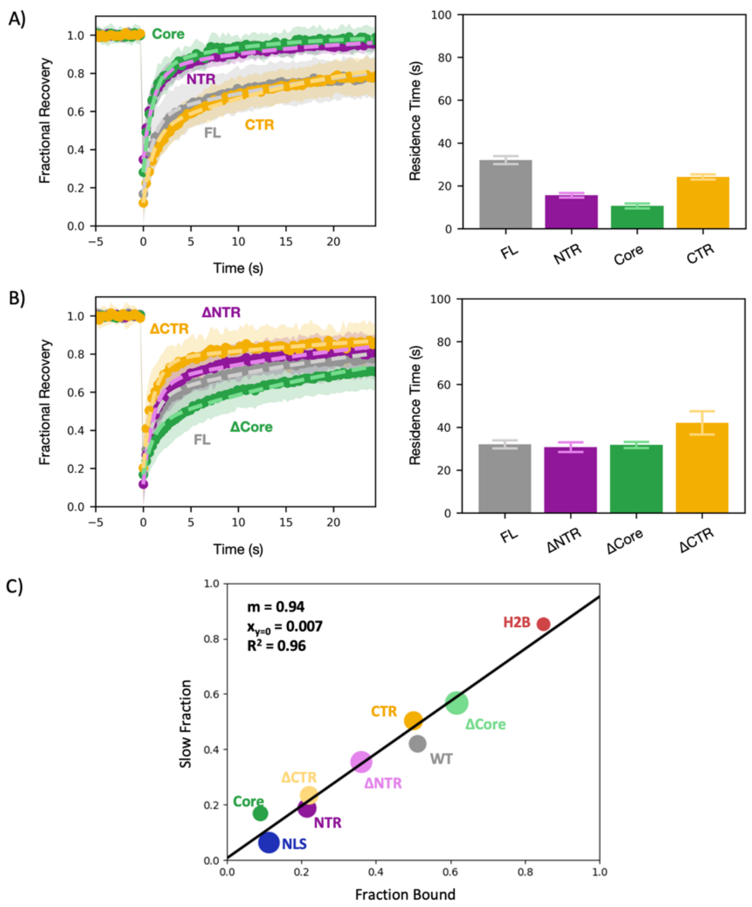


**Figure S3—FRAP confirms SMT findings.**

(A) FRAP plots of p300 regions (left) with residence times plotted (right). (B) FRAP plots of p300 truncations (left) with residence times plotted (right). (C) Scatter plot of fraction bound (SMT) and slow fraction (FRAP) for transgene constructs shows a high degree of correlation. The size of each point encompasses both the standard deviation of the bound fraction and the 95% confidence interval of the slow fraction.

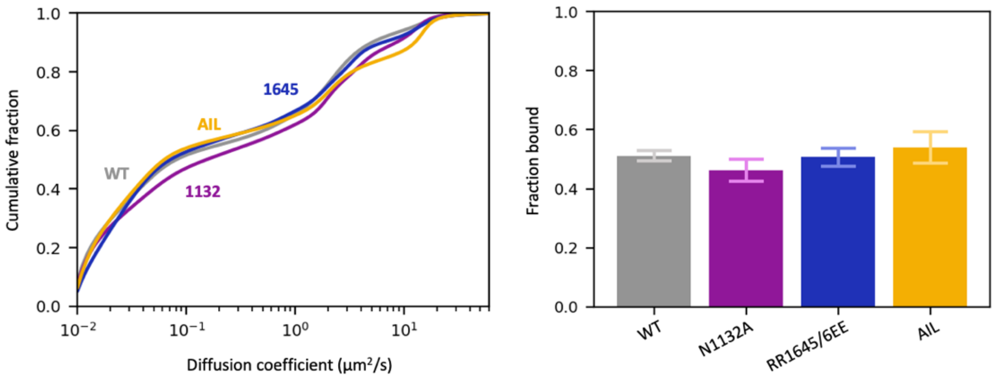


**Figure S4—SMT of Core mutants.**
SMT plots as in Figure 1 of Core mutants shown in vitro to have large effects on p300 catalytic activity: N1132A [S1], RR1645/6EE [S2], autoinhibitory loop $[K>R]_9$ (AIL) [S3].

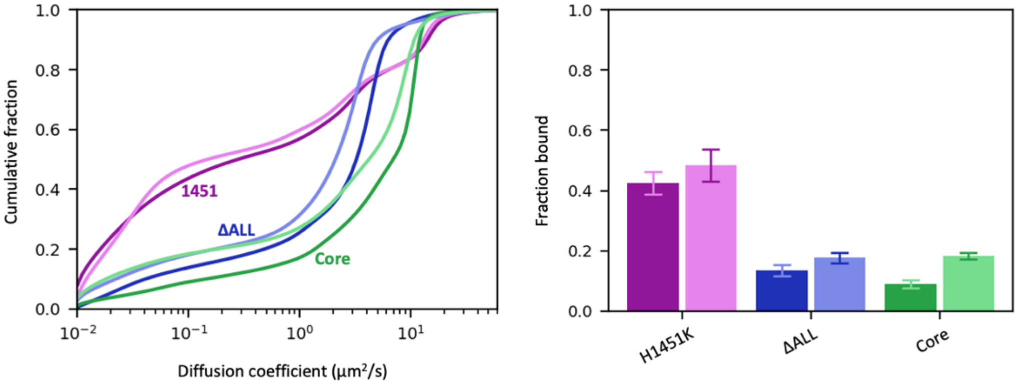


**Figure S5—SMT of Core mutants**
SMT plots as in Figure 4 of mutants ± A485: the inhibitor-resistant mutant 1451 has no significant change, while the two unbound constructs ΔAll and Core show mild increases upon addition of the inhibitor.

### SUPPLEMENTAL TABLES

| Sample | Mean | SD | Num. cells | $D_{unbound}$ | # Trajs. | Figure |
|---|---|---|---|---|---|---|
| H2B | 0.847 | 0.010 | 26 | 6.1 | 76,546 | 1 |
| Halo-NLS | 0.110 | 0.013 | 45 | 17.1 | 9,836 | 1 |
| Halo-p300 KI-1 | 0.551 | 0.014 | 32 | 2.9 | 26,951 | 1 |
| Halo-p300 KI-2 | 0.521 | 0.017 | 33 | 2.7 | 34,664 | 1 |
| WT | 0.510 | 0.020 | 25 | 2.2 | 19,200 | 1 |
| Core | 0.087 | 0.009 | 30 | 11.8 | 15,732 | 2 |
| CTR | 0.502 | 0.022 | 36 | 6.7 | 21,276 | 2 |
| NTR | 0.227 | 0.014 | 29 | 7.4 | 31,228 | 2 |
| ΔCore | 0.616 | 0.024 | 30 | 3.5 | 15,613 | 2 |
| ΔCTR | 0.221 | 0.019 | 31 | 5.6 | 12,339 | 2 |
| ΔNTR | 0.359 | 0.017 | 30 | 4.2 | 11,499 | 2 |
| IBID | 0.174 | 0.015 | 39 | 14.2 | 17,464 | 3 |
| NRID | 0.105 | 0.009 | 40 | 14.2 | 27,129 | 3 |
| TAZ1 | 0.192 | 0.014 | 40 | 15.6 | 23,394 | 3 |
| TAZ2 | 0.236 | 0.021 | 39 | 14.2 | 14,974 | 3 |
| ΔALL | 0.132 | 0.010 | 40 | 4.6 | 10,590 | 3 |
| ΔIBID | 0.411 | 0.017 | 25 | 3.9 | 10,344 | 3 |
| ΔKIX | 0.331 | 0.028 | 30 | 2.9 | 18,565 | 3 |
| ΔNRID | 0.539 | 0.037 | 20 | 2.2 | 6,322 | 3 |
| ΔTAZ1 | 0.326 | 0.031 | 21 | 2.4 | 6,028 | 3 |
| ΔTAZ2 | 0.430 | 0.026 | 29 | 3.9 | 30,821 | 3 |
| 1451 | 0.426 | 0.042 | 15 | 15.6 | 6,698 | 4 |
| 1467 | 0.602 | 0.019 | 65 | 2.4 | 13,938 | 4 |
| 1451+A485 | 0.487 | 0.048 | 19 | 14.2 | 8,185 | 4 |
| NTR-Core+A485 | 0.252 | 0.020 | 24 | 15.6 | 8,431 | 4 |
| WT+A485 | 0.587 | 0.023 | 30 | 2.0 | 6,639 | 4 |
| ΔIBID+A485 | 0.431 | 0.021 | 42 | 2.2 | 10,601 | 4 |
| ΔKIX+A485 | 0.533 | 0.019 | 41 | 2.0 | 13,995 | 4 |
| ΔNRID+A485 | 0.725 | 0.022 | 41 | 1.3 | 13,817 | 4 |
| ΔNTR +A485 | 0.500 | 0.026 | 32 | 3.5 | 19,688 | 4 |
| ΔTAZ1+A485 | 0.515 | 0.023 | 41 | 1.8 | 15,377 | 4 |
| ΔTAZ2+A485 | 0.489 | 0.027 | 41 | 2.9 | 14,598 | 4 |
| 1132 | 0.468 | 0.022 | 51 | 2.0 | 20,074 | S |
| 1645 | 0.517 | 0.021 | 43 | 3.5 | 11,325 | S |
| AIL | 0.532 | 0.037 | 40 | 15.6 | 18,954 | S |
| Core+A485 | 0.181 | 0.013 | 19 | 8.9 | 5,439 | S |
| ΔALL+A485 | 0.174 | 0.014 | 20 | 3.5 | 7,646 | S |

**Table S1:** Summary of SMT analysis for all constructs. $D_{unbound}$ is the diffusion coefficient for the predominant population >1 $\mu m^2/s$. Number of trajectories refers to the total of the whole set.

| Sample | Mean Fluorescence | Est. # molecules | Figure |
|---|---|---|---|
| Halo-NLS | 39222 | 2,309,500 | 1 |
| Halo-p300 KI-1 | 1082 | 49,000 | 1 |
| Halo-p300 KI-2 | 2437 | 129,300 | 1 |
| WT | 324 | 4,100 | 1 |
| Core | 700 | 26,400 | 2 |
| CTR | 656 | 23,800 | 2 |
| NTR | 840 | 34,700 | 2 |
| ΔCore | 522 | 15,800 | 2 |
| ΔCTR | 532 | 16,400 | 2 |
| ΔKIX | 601 | 20,500 | 2 |
| ΔNTR | 603 | 20,600 | 2 |
| IBID | 1407 | 68,300 | 3 |
| KIX | 3068 | 166,700 | 3 |
| NRID | 9710 | 560,400 | 3 |
| TAZ1 | 2975 | 161,200 | 3 |
| TAZ2 | 850 | 35,300 | 3 |
| ΔALL | 686 | 25,500 | 3 |
| ΔIBID | 226 | NC | 3 |
| ΔNRID | 388 | 7,900 | 3 |
| ΔTAZ1 | 417 | 9,600 | 3 |
| ΔTAZ2 | 533 | 16,500 | 3 |
| 1451 | 461 | 12,200 | 4 |
| 1467 | 692 | 25,900 | 4 |
| CTCF | 2112 | 110,000 | S2 |
| Negative control | 256 | 0 (fixed) | S2 |
| 1132 | 459 | 12,100 | S4 |
| 1645 | 513 | 15,300 | S4 |
| AIL | 510 | 15,100 | S4 |

**Table S2:** Summary of flow cytometry in Fig. S5. Number of molecules are estimated according to a previously published protocol [S4]. NC: not computed.

| Sample | *A* | *B* | $k_a$ | $k_b$ | $R^2$ | RMSE | # Cells |
|---|---|---|---|---|---|---|---|
| Halo NLS | 0.32 ± 0.01 | 0.61 | 1.75 ± 0.12 | 0.10 ± 0.01 | 0.989 | 0.006 | 42 |
| H2B | 0.010 ± 0.001 | 0.13 | 0.93 ± 0.31 | $1\times10^{-4}$ ± $1\times10^{-4}$ | 0.736 | 0.001 | 18 |
| KI-1 | 0.36 ± 0.01 | 0.28 | 0.95 | 0.046 ± 0.002 | 0.988 | 0.012 | 29 |
| KI-2 | 0.35 ± 0.01 | 0.27 | 0.95 | 0.038 ± 0.002 | 0.983 | 0.015 | 30 |
| WT | 0.41 ± 0.01 | 0.17 | 0.95 | 0.031 ± 0.002 | 0.983 | 0.015 | 29 |
| NTR | 0.46 ± 0.01 | 0.35 | 0.95 | 0.064 ± 0.005 | 0.989 | 0.011 | 30 |
| Core | 0.55 ± 0.02 | 0.28 | 0.95 | 0.093 ± 0.011 | 0.982 | 0.016 | 26 |
| CTR | 0.38 ± 0.01 | 0.12 | 0.95 | 0.041 ± 0.002 | 0.983 | 0.018 | 30 |
| ΔNTR | 0.53 ± 0.01 | 0.12 | 0.95 | 0.032 ± 0.003 | 0.983 | 0.017 | 36 |
| ΔCore | 0.26 ± 0.01 | 0.17 | 0.95 | 0.031 ± 0.001 | 0.984 | 0.015 | 29 |
| ΔCTR | 0.56 ± 0.01 | 0.20 | 0.95 | 0.024 ± 0.004 | 0.975 | 0.017 | 23 |
| WT + A485 | 0.32 ± 0.01 | 0.12 | 0.95 | 0.027 ± 0.001 | 0.985 | 0.014 | 40 |
| H1451K | 0.42 ± 0.01 | 0.14 | 0.95 | 0.029 ± 0.001 | 0.984 | 0.015 | 39 |
| H1451K + A485 | 0.48 ± 0.02 | 0.03 | 0.95 | 0.029 ± 0.002 | 0.971 | 0.024 | 37 |

**Table S3:** Fitting parameters with 95% confidence intervals from FRAP allowing *A*, $k_a$, to vary for all constructs. The values of $k_b$ were fixed for all p300 variants as described in the methods.

**Supplemental Note**

We observed that the CTR fragment, comprising the TFIDs TAZ2 and IBID, has a higher fraction bound than either TFID alone. One possible explanation for this, discussed in the main text, is that both TFIDs simultaneously engage adjacent transcription factors clustered at adjacent sites on chromatin, resulting in an increase in *strength* of interaction. An alternative model is that the TFIDs bind independently to different TF-bound sites, and combining the two TFIDs increases the fraction bound by increasing the *number* of interactions. We argue here that the latter model disagrees quantitatively with our results.

Suppose that a protein with unbound concentration $A$ can bind multiple types of sites on chromatin where the concentration of site $i$ is given by $B_i$. Let the concentration of the protein-DNA complex $i$ be $C_i$. Suppose that the protein binds only one site at a time.

Because our experiments have shown that the bound fraction of p300 and its mutants is insensitive to the protein expression level, we assume that binding sites are in excess ($B_i \gg A$) and that the concentration of unbound sites of type $i$ is approximately equal to $B_i$. In this limit, the dissociation constant of the $i^{\text{th}}$ type of site is given by

$$K_{di} = \frac{AB_i}{C_i} \tag{1}$$

The overall fraction bound (to any site) will be given by

$$f_b = \frac{\sum_i C_i}{A + \sum_i C_i} = \frac{\sum_i x_i}{1 + \sum_i x_i} \tag{2}$$

where we have defined

$$x_i \equiv \frac{C_i}{A} = \frac{B_i}{K_{di}} \tag{3}$$

This quantity, $x_i$, is essentially the concentration of the $i^{\text{th}}$ type of binding site in units of its $K_{di}$.

Suppose that we have a series of mutants of the protein that can each bind only a single type of site. The fraction bound of the mutant that binds only site $j$ will be

$$f_{b,j} = \frac{C_j}{A + C_j} = \frac{x_j}{1 + x_j} \tag{4}$$

Rearranging gives

$$x_j = \frac{f_{b,j}}{1 - f_{b,j}} \tag{5}$$

Substituting into equation (6) gives

$$f_b = \frac{\sum_i \frac{f_{b,i}}{1-f_{b,i}}}{1 + \sum_i \frac{f_{b,i}}{1-f_{b,i}}} \tag{6}$$

We can thus predict the bound fraction of the wild-type protein from the bound fractions of mutants that bind disjoint subsets of sites.

Estimating the fraction bound of the full CTR from the fractions bound of TAZ2 and IBID gives

$$f_b = \frac{\frac{0.23}{1-0.23} + \frac{0.17}{1-0.17}}{1 + \frac{0.23}{1-0.23} + \frac{0.17}{1-0.17}} = 0.33 \tag{7}$$

While this is greater than either TFID alone, it is substantially less than the observed bound fraction of 0.50, suggesting that independent binding of the two TFIDs to distinct sites is not sufficient to account for the observed increase in bound fraction.
